# Expanding tunable selection in yeast using auxotrophic markers *URA3* and *TRP1*

**DOI:** 10.1101/2025.08.19.671079

**Authors:** Yunwei Luo, Tatsuhisa Tsuboi

## Abstract

Auxotrophic markers are fundamental tools in yeast genetics, but their use is traditionally binary (growth or no growth), which limits quantitative analysis. Here we repurpose the classic auxotrophic markers *URA3* and *TRP1* as tunable selection tools that quantitatively link gene-expression levels to growth outcomes. In uracil-deficient medium (SC–Ura), *URA3* yields a positive, semi-quantitative growth response proportional to its expression level. Conversely, under selection with 5-fluoroanthranilic acid (5-FAA), *TRP1* shows an inverse relationship between expression and survival: higher *TRP1* expression triggers greater 5-FAA toxicity and reduced growth. Using integrative plasmids to vary gene copy number, we tuned expression levels and found that even ∼2–4-fold differences produce distinct growth phenotypes. We also obtained dose–response results correlating marker expression with growth rates. Finally, applying this system to a synthetic mitochondrial mRNA localization scheme, we detected the resulting increase in protein expression via the growth-based readout. By converting traditional binary markers into tunable, quantitative selectors, this platform expands the yeast toolkit for evaluating gene expression differences and synthetic circuit function with simple growth assays.

## Introduction

Classical yeast genetic selections rely on auxotrophic marker genes (e.g., *HIS3, URA3, TRP1, LEU2*) to confer prototrophy versus auxotrophy in a binary fashion (growth vs. no growth). These markers typically yield all-or-none outcomes unless modified by additional strategies. In particular, *HIS3* has long been paired with 3-aminotriazole (3-AT) to achieve graded selection: 3-AT is a competitive inhibitor of the *HIS3*-encoded imidazoleglycerol-phosphate dehydratase, so increasing 3-AT in SC–His raises the *HIS3* activity required for growth ^1^. This “titration” strategy is widely used in yeast two-hybrid and promoter-strength assays to fine-tune reporter stringency and reduce background growth^1^. For example, higher 3-AT levels demand higher *HIS3* expression to overcome inhibition, thereby eliminating false positives in protein–protein interaction screens. However, 3-AT is specific to the histidine pathway and the *HIS3* product, limiting applicability beyond *HIS3*-based reporters. Historically, other markers have lacked analogous tunable regulators, so selections using *URA3* or *TRP1* have remained essentially binary.

The *URA3* gene encodes orotidine-5′-phosphate decarboxylase (ODCase), a key enzyme in pyrimidine biosynthesis. In the presence of the toxic analog 5-fluoroorotic acid (5-FOA), ODCase converts 5-FOA into 5-fluorouracil, killing *URA3*^+^ cells (Figure 1A)^2^. Thus, 5-FOA has been exploited for negative selection: only Ura^−^ cells grow, making 5-FOA useful for selecting *URA3* loss. Similarly, *TRP1* (phosphoribosylanthranilate isomerase in tryptophan biosynthesis) can be counterselected using 5-fluoroanthranilic acid (5-FAA). Cells with functional *TRP1* convert 5-FAA (an anthranilate analog) into toxic tryptophan analogs (e.g., 5-fluorotryptophan or 5-methyltryptophan), causing cell death (Figure 1B)^3^. Despite these developments, *URA3* and *TRP1* have traditionally been used only in binary modes—positive selection on dropout media or negative selection on 5-FOA/5-FAA—rather than as tunable, gene-dosage-dependent reporters.

**Figure 1.**
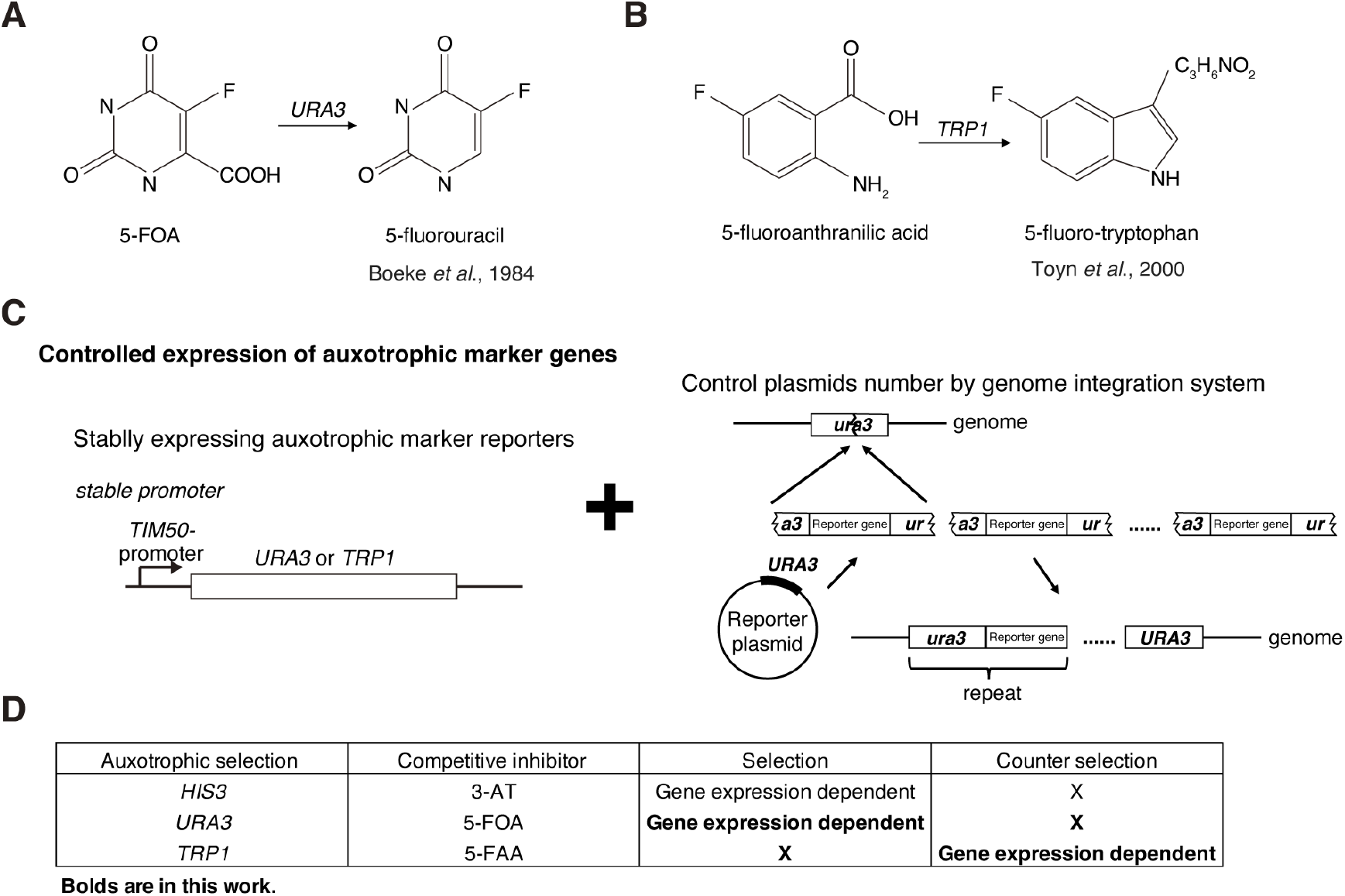
Expanding tunable selection tools in yeast using *URA3* and *TRP1* markers. (A) *URA3* converts 5-FOA to 5-fluorouracil, leading to the accumulation of fluorinated uracil analogs that inhibit DNA synthesis. (B) *TRP1* converts 5-FAA to 5-fluorotryptophan (and other toxic tryptophan analogs), which disrupts protein synthesis. (C) Schematic of the tunable selection approach. Left: *URA3* and *TRP1* are expressed under the modest *TIM50* promoter. Right: Multiple copies of the integrative plasmid (carrying *URA3* or *TRP1*) can insert into the genome, resulting in different expression levels of the marker gene. (D) Complementary selection regimes for the two markers. *URA3* on SC–Ura supports growth proportional to expression level (higher expression = better growth, positive selection), whereas *TRP1* under 5-FAA yields growth inversely proportional to expression (higher expression = more toxic conversion, less growth, negative selection).

Here, we introduce a strategy to convert *URA3* and *TRP1* into tunable reporters of expression level. We show that *URA3* selection systems, analogous to *HIS3*/3-AT, yield growth phenotypes concordant with gene expression level, whereas *TRP1* under 5-FAA yields growth phenotypes inverse to expression level. In other words, this dual-marker design provides a bidirectional selection: *URA3* acts as a positive, semi-quantitative selector, and *TRP1*/5-FAA acts as a negative, expression-inverted selector. We construct yeast strains with controlled *URA3* or *TRP1* expression using the moderate *TIM50* promoter and vary marker dosage by adjusting the copy number of integrated plasmids. We examine growth on SC–Ura and SC–Trp as a function of marker dosage, as well as on counterselective SC media containing 5-FOA or 5-FAA. In addition, to quantitatively probe this system, we employ a model in which mRNA localization to mitochondria increases protein production by coupling translation with protein import and potentially improving folding efficiency ^4^. Recent studies show that mitochondrial mRNA localization is both necessary and sufficient to boost production of certain proteins under respiratory conditions^5^. We find that bringing an mRNA into close proximity to mitochondria enhances its translation, which alters growth on selective media compared with an unlocalized control. Our results reveal clear gene dose–dependent growth effects: increased expression enhances growth on dropout media for *URA3*, but exacerbates drug sensitivity under *TRP1* counterselection. Collectively, this work establishes a new quantitative selection platform in yeast that links marker gene dosage to a convenient growth readout, enabling evaluation of promoter strength and gene expression in yeast and providing a valuable tool for synthetic biology and gene circuit engineering.

## Results

### Tuning auxotrophic marker expression

To validate the use of auxotrophic markers as reporters of relative protein expression, we first confirmed that varying the expression levels of *URA3* or *TRP1* produces distinguishable growth phenotypes. Quantitative transcript measurements in yeast indicate a median mRNA abundance of ∼5–10 molecules per cell ^5^. *URA3* and *TRP1* are typically expressed at ∼7–9 mRNA molecules per cell ^5^. Therefore, we selected the *TIM50* promoter—which is a moderate constitutive promoter yielding ∼4–6 transcripts per cell ^4,5^ —to drive expression of the marker genes (Figure 1C, left). We also used a yeast integrative plasmid system to generate isogenic strains carrying different genomic copy numbers of the same marker construct (Figure 1C, right). All strains derive from the W303-1A background (*ura3*^−^, *trp1*^−^) and thus require functional *URA3* or *TRP1* transgenes for prototrophic growth. In this study, we introduce and characterize *URA3* selection behaves analogously to *HIS3*/3-AT with graded output, and *TRP1* under 5-FAA provides an inverse readout: *URA3* shows growth phenotypes concordant with expression level, whereas *TRP1* under 5-FAA counter-selection shows growth phenotypes inverse to expression level (Figure 1D).

### Tuning *URA3* expression correlates with auxotrophic growth

When strains were spotted onto synthetic complete plates lacking uracil (SC–Ura), those with higher *URA3* copy number grew more robustly than the 1-copy strain (Figure 2A, B). The 4-copy *URA3* strain formed notably larger colonies and grew faster than the 1-copy strain, suggesting that a single *URA3* gene under the *TIM50* promoter provides suboptimal complementation of the *ura3* auxotrophy, whereas multiple copies significantly improve growth. We next evaluated 5-FOA counterselection as a potentially more sensitive readout of expression differences by plating the *URA3* copy-number series on media containing graded concentrations of 5-FOA (Figure 2B). As expected, all *URA3*-expressing strains were inhibited on 5-FOA, since Ura3p converts 5-FOA into a toxic metabolite (Figure 1A). At a standard concentration of 1000 µg/mL, a single *URA3* copy substantially impaired growth. Reducing the 5-FOA dose down to 300 µg/mL did not further differentiate the strains—lower *URA3* copy number did not mitigate sensitivity beyond an already lethal baseline. At these doses, none of the *URA3*^+^ strains were able to form colonies. Thus, the traditional *URA3*/5-FOA system is highly stringent: once any functional Ura3p is present (even at one gene copy), 5-FOA imposes such strong negative selection that it obscures gradations in expression. Taken together, these observations show that *URA3* expression level can be assessed in a semi-quantitative manner by growth on uracil-dropout plates (higher plasmid copy yields a clear growth advantage), whereas 5-FOA in the 300–1000 µg/mL range does not provide a gradable window.

**Figure 2.**
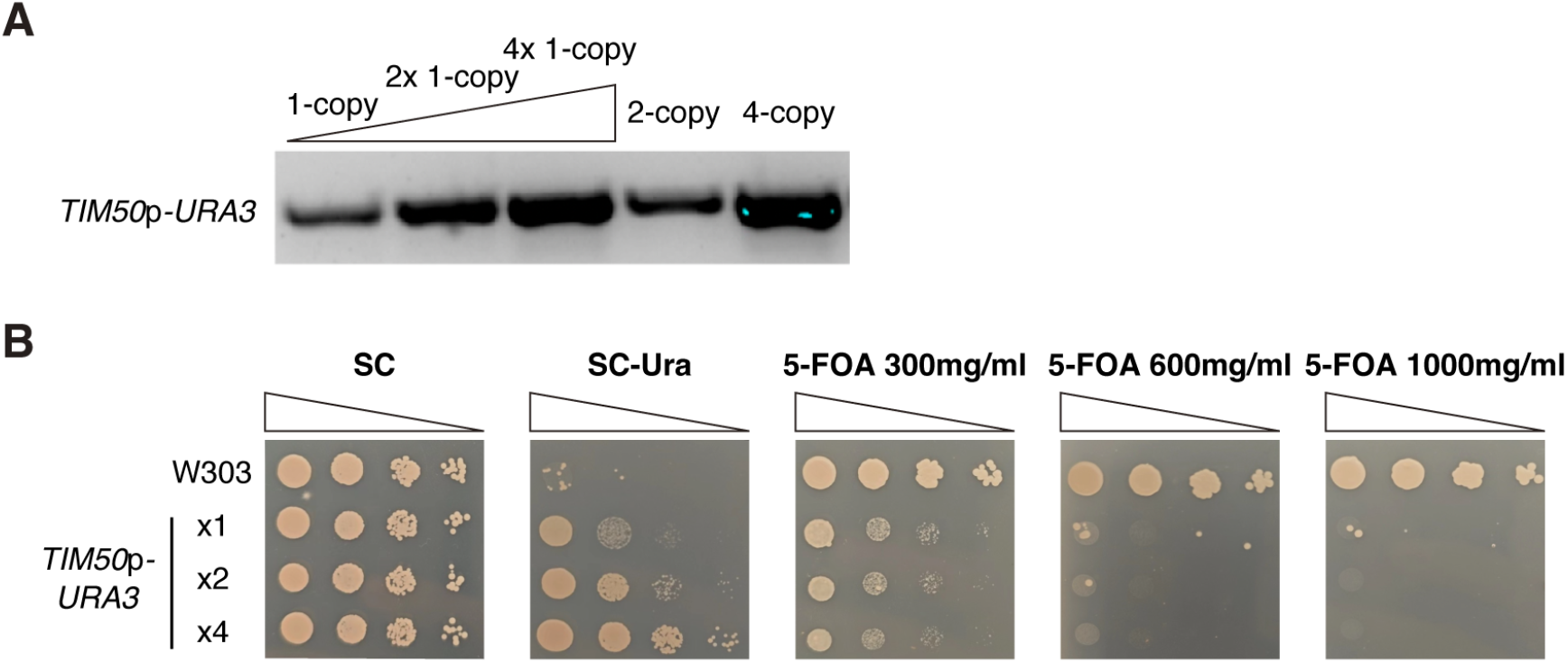
*URA3* copy number correlates with growth on uracil-dropout medium. (A) PCR confirmation of *URA3* copy number. Gel electrophoresis of PCR products was used to verify 1-copy, 2-copy, and 4-copy *URA3* integrants. (B) Spotting assay of strains with different *URA3* copy numbers on SC–Ura plates and 5-FOA plates (300–1000 µg/mL 5-FOA) after 2 days of inoculation at 30 °C.

### *TRP1* expression inversely correlates with 5-FAA counterselection

In contrast to *URA3*, increasing *TRP1* copy number did not produce a marked growth benefit on tryptophan-dropout medium (SC–Trp; Figure 3A,B). Even a single *TRP1* copy was sufficient to sustain near wild-type growth, suggesting that Trp1p activity was not strongly growth-limiting under these conditions. We next evaluated 5-fluoroanthranilic acid (5-FAA) as a more sensitive readout of expression differences. Under initial “standard” counterselection conditions (SC base with 5% glucose, 500 µg/mL 5-FAA, and only a trace of tryptophan to permit minimal growth^3^), both the 1-copy and 2-copy *TRP1* strains failed to grow. This outcome mirrored the high stringency observed for *URA3* on 5-FOA. We therefore fine-tuned the conditions by lowering the 5-FAA concentration and adding a minute amount of tryptophan to identify a regime that allows intermediate growth. Across a series of 5-FAA and nutrient titrations, we found conditions that produced partial growth for *TRP1*-expressing cells. Ultimately, we identified an optimized 5-FAA concentration (with 0.001% tryptophan to avoid complete starvation ^3^) that yielded a reproducible growth window (Figure 3B). Under these conditions, the 1-copy *TRP1* strain grew slightly better than the 2-copy strain, consistent with the idea that higher Trp1p levels generate more toxic fluorotryptophan from 5-FAA. Together, these results establish that the *TRP1*/5-FAA system can be configured as an anti-expression selection: growth decreases as *TRP1* expression increases, provided the counterselection window is properly tuned.

**Figure 3.**
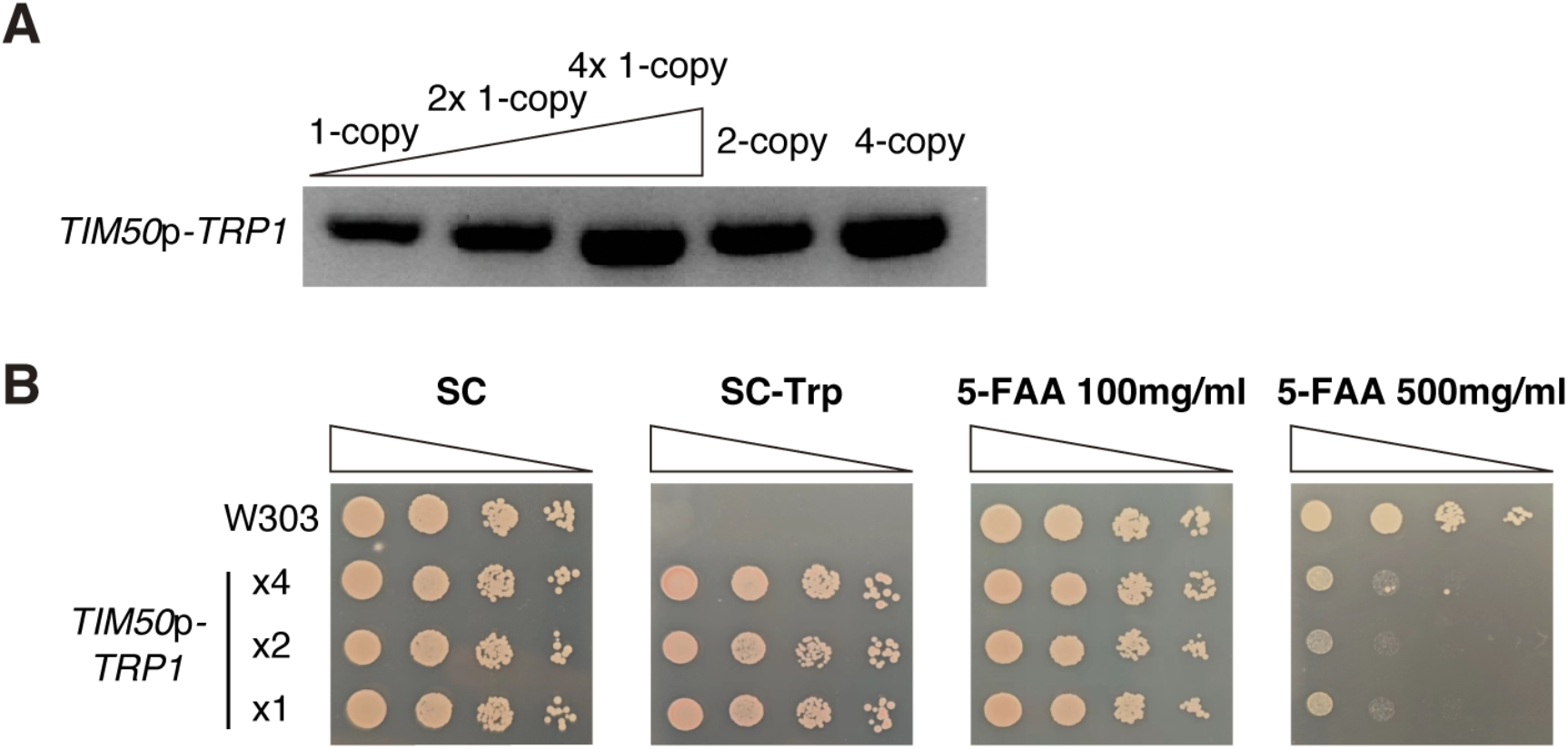
Tuning *TRP1* expression with tryptophan dropout and calibrated 5-FAA selection. (A) PCR confirmation of *TRP1* copy number. Gel electrophoresis of PCR products was used to verify 1-copy, 2-copy, and 4-copy *TRP1* integrants. (B) Spotting assay of strains with different *TRP1* copy numbers on SC–Trp plates and under optimized 5-FAA counterselection (with 0.001% tryptophan to avoid complete starvation) after 2 days of inoculation at 30 °C.

### Artificial tethering of mRNA to mitochondria alters 5-FAA sensitivity of *TRP1*-expressing cells

To test whether localizing an mRNA to the mitochondrial surface alters translation enough to impact growth, we employed the MS2–MCP tethering system. We appended 12× MS2 stem-loop sequences to the 3′ UTR of the *URA3* or *TRP1* reporter gene. We also genomically integrated a fusion of the MS2 coat protein (MCP) to the outer-membrane protein Tom70 under a constitutive promoter. In this configuration, MCP is anchored on the cytosolic face of mitochondria and binds the MS2-tagged mRNAs, recruiting them to the organelle surface (Figure 4A, B). In this configuration, the tethered reporter showed an ∼10-fold increase in protein expression^4^. Baseline growth on rich media or simple dropout media was unchanged by the presence of Tom70–MCP. However, under selective conditions for *TRP1*, tethering the mRNA produced a marked shift in 5-FAA sensitivity (Figure 4C). Under the optimized 5-FAA regime (2% glucose plates with tuned 5-FAA dose), the tethered *TRP1* strain exhibited much stronger growth inhibition—forming only tiny colonies or none at all—whereas the untethered control formed readily visible colonies. Thus, mitochondrial localization of *TRP1* mRNA increases effective Trp1p activity, enhancing conversion of 5-FAA to toxic products and heightening drug sensitivity. By contrast, for *URA3*, tethering did not detectably change 5-FOA sensitivity: all *URA3*^+^ cells remained fully 5-FOA– sensitive, consistent with the uniformly high stringency of *URA3*/5-FOA selection (Figure 4D). As expected from auxotrophic complementation, growth on SC–Ura still reflected expression level (excess Ura3p improved growth), whereas 5-FOA counterselection was too strict to reveal any subtle changes due to tethering.

**Figure 4.**
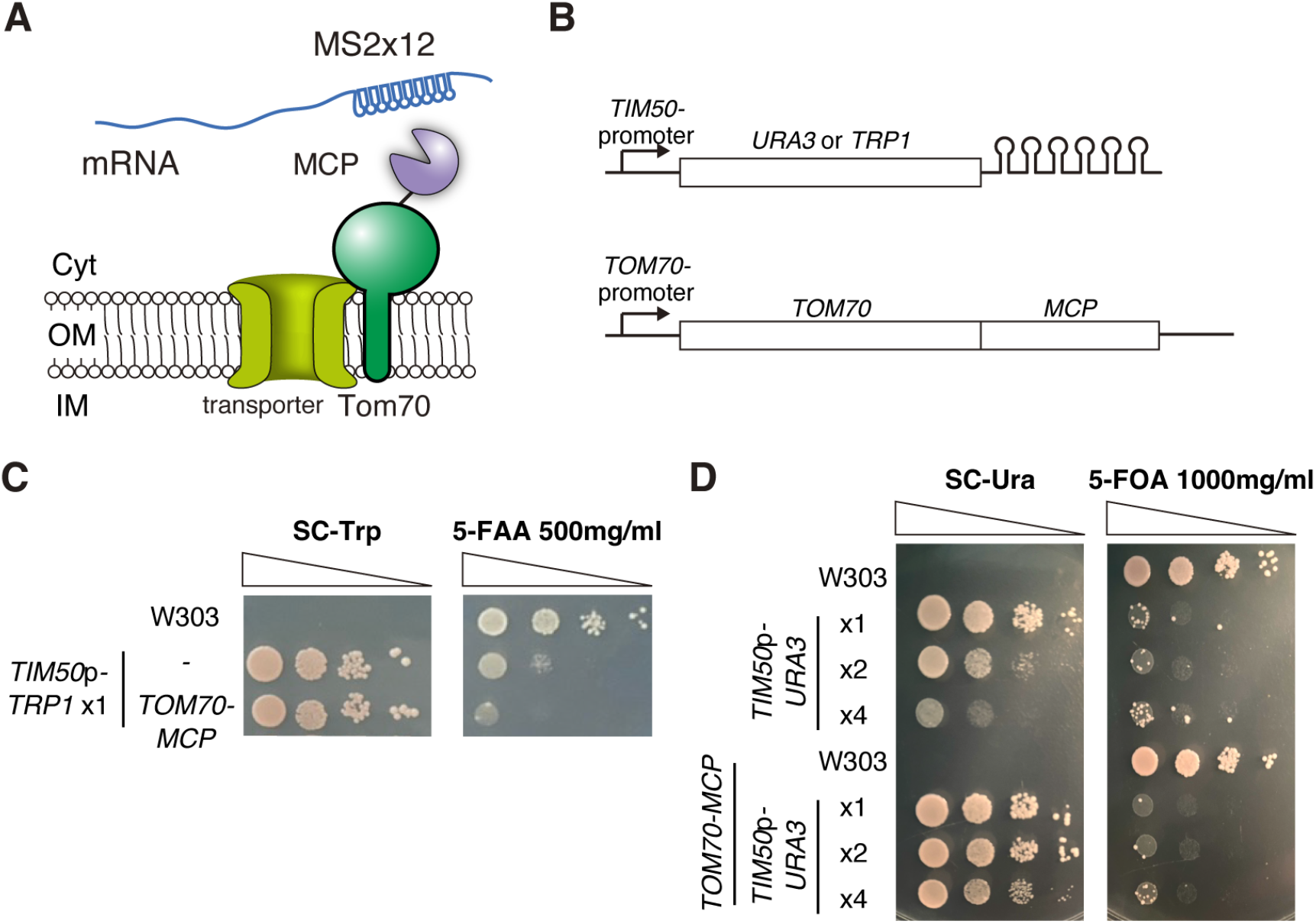
MS2–MCP-mediated mitochondrial tethering enhances *TRP1* toxicity under 5-FAA selection. (A) Schematic of the MS2–MCP tethering system for mRNA localization. MCP (MS2 coat protein) is fused to Tom70 and anchored on the mitochondrial outer membrane, where it binds MS2 hairpin tags on the target mRNA, localizing the mRNA to the mitochondrial surface. (B) Diagram of strain construction for artificial mRNA localization. The TOM70–MCP fusion is genomically integrated into *URA3*-MS2 or *TRP1*-MS2 strains, allowing the *URA3* or *TRP1* mRNA to be tethered to mitochondria. Both *URA3* and *TRP1* are expressed at low copy number under the *TIM50* promoter in these strains. (C) Spotting assay of *TRP1* strains with and without tethering on SC–Trp plates and 5-FAA plates (with 2% glucose and 0.001% tryptophan to avoid complete starvation) after 2 days of inoculation at 30 °C. (D) Spotting assay of *URA3* strains with and without tethering under tuned SC-URA and 5-FOA conditions after 2 days of inoculation at 30 °C.

## Discussion

This study presents a simple, tunable framework that converts differences in gene expression into predictable growth outcomes using classic auxotrophic markers. By driving *URA3* or *TRP1* from a modest promoter (*TIM50*) and constructing isogenic copy-number series, we resolved two complementary regimes—essentially a bidirectional selection approach: a pro-expression selection in which growth scales with marker abundance (*URA3* on SC–Ura) and an anti-expression selection in which growth inversely tracks marker abundance (*TRP1* under 5-FAA counterselection, after calibration). Together, these regimes transform auxotrophic markers from binary selectors into quantitative meters with configurable dynamic range. Three design principles underlie this behavior. (i) Promoter strength and copy number define the measurable window: *URA3* driven by *TIM50* promoter behaves as a semi-quantitative rheostat on SC–Ura (a single copy is suboptimal, and additional copies improve growth), whereas *TRP1* saturates on SC–Trp with a single copy. (ii) Counterselection chemistry sets the ceiling: along the *URA3*/5-FOA axis, once any Ura3p is present, 5-FOA (300–1000 µg/mL) collapses phenotypes to an on/off response. (iii) Nutrient and inhibitor titration can linearize otherwise binary responses: lowering 5-FAA and adding trace tryptophan (0.001%) reveals a stable window in which growth decreases as *TRP1* expression increases. This graded, expression-inverted regime mirrors tunable counterselections such as *HIS3*/3-AT, with the midpoint set by the 5-FAA dose and nutrient context.

The distinct behaviors of *URA3* and *TRP1* can be explained by both toxic metabolite chemistry and biosynthetic pathway positioning. For *URA3*, the toxic intermediate generated from 5-FOA creates a threshold effect: even minimal Ura3p activity is sufficient to convert 5-FOA into 5-fluorouracil (5-FU), a pyrimidine analog that is incorporated into RNA and DNA, disrupting nucleic acid processing and rapidly killing cells ^6^. Thus, once *URA3* is expressed at any detectable level, 5-FOA imposes strong negative selection, and additional expression offers little further discrimination. Pathway structure reinforces this outcome. *URA3* encodes the final enzyme in uracil biosynthesis, decarboxylating orotidine-5′-phosphate to uridine monophosphate (UMP). Precursor synthesis by upstream enzymes (*URA2, URA4*, and others) is typically not limiting, meaning that Ura3p sets the decisive bottleneck in uracil production^7^. Accordingly, *URA3* expression intensity directly determines intracellular UTP levels, influencing transcriptional capacity and ultimately proliferation. This explains why *URA3* dosage produces graded growth differences on SC– Ura, while FOA selection collapses to a binary phenotype. In contrast, *TRP1* encodes phosphoribosylanthranilate isomerase, an intermediate enzyme in tryptophan biosynthesis. Downstream enzymes (Trp3p, Trp4p, and Trp5p) control the conversion of indole intermediates into tryptophan^8^. As a result, varying *TRP1* dosage has little impact under tryptophan dropout: even at the lowest *TRP1* copy number, sufficient precursor is available for the downstream enzymes to support near-normal growth. Only under 5-FAA counterselection does *TRP1* dosage become informative. 5-FAA acts both as a substrate for Trp1p, generating toxic fluorotryptophan analogs, and as a feedback inhibitor of anthranilate synthase, the first enzyme in the pathway. This dual mechanism amplifies the growth penalty of higher *TRP1* expression while simultaneously limiting tryptophan biosynthesis, producing a tunable, expression-inverted response when drug and nutrient conditions are calibrated. Together, these insights show how differences in toxic metabolite metabolism and pathway bottlenecks explain the contrasting behaviors of *URA3* and *TRP1*. In principle, the same calibration logic could extend to other auxotrophic markers (e.g., *LEU2, ADE2, LYS2*) and their analogs, provided their drug chemistries yield a resolvable window.

A key functional test demonstrated that the anti-expression window is sensitive to where translation occurs, not just how much transcript is produced. Tethering *TRP1* mRNA to the mitochondrial surface via MS2– MCP (Tom70–MCP) caused a marked leftward shift in the 5-FAA growth response. Under the optimized conditions, the tethered strain formed only minute or no colonies, compared to visible colonies in the untethered controls. This effect is consistent with enhanced effective *TRP1* activity resulting from mitochondrial localization of the mRNA (and consequently the protein), and it demonstrates that the *TRP1*/5-FAA window can report local translation gains driven by subcellular targeting. By contrast, the *URA3*/5-FOA axis remained uniformly stringent regardless of tethering, reinforcing its limited utility for fine discrimination of expression changes. Together, these observations indicate that the *TRP1*/5-FAA configuration can serve as a sensitive sensor for perturbations that amplify translation at organelle surfaces. For example, candidate factors that could be screened with this system include RNA-binding proteins (e.g., Puf3p, which binds many mitochondria-localized mRNAs ^9^), components of the TOM complex (notably, mRNA localization is reduced upon TOM20 deletion ^10^), and general translation factors. Prior work showing that slowed translation elongation increases mitochondrial mRNA localization and can boost protein output ^4^ supports a model in which ribosome kinetics and organelle-proximal factors jointly tune localized translation. In this context, suppressor mutations may relieve a translational pause or dependency created by enforced tethering, providing a potential avenue to identify new factors that influence localized translation.

Limitations of our framework point to straightforward refinements. In our hands, the *URA3*/5-FOA window remained binary across 300–1000 µg/mL 5-FOA. We note that genetic background can modulate this stringency—e.g., overexpression of YJL055W confers resistance to 5-FOA/5-FU ^6^—suggesting that engineered resistance alleles could broaden the *URA3* counterselection window. Similarly, the *TRP1*/5-FAA window is media-dependent, necessitating calibration of carbon source and tryptophan levels in each experimental context to avoid false binary outcomes. Additionally, the dynamic range and midpoint of the selection will shift with promoter strength and genomic context. While using *TIM50*-driven integrants provided a controlled baseline here, alternate expression architectures will require re-titration of selection conditions. Finally, extending plate-based readouts to liquid culture growth curves would enable continuous dose–response fitting and sharper quantitative comparisons.

In sum, by configuring *URA3* for positive selection on dropout media and *TRP1* for calibrated counterselection with 5-FAA, we obtain a compact, inexpensive toolkit that reports gene expression levels—and, critically, subcellular translation enhancements—through simple growth assays. The ability of mitochondrial mRNA tethering to sharpen 5-FAA sensitivity underscores the framework’s utility for detecting translation-enhancing elements (such as localization signals or other translational regulators). Furthermore, this dual-selection approach can streamline the rapid prototyping of expression cassettes, gene circuits, and localization strategies in yeast synthetic biology by converting molecular design choices into differential survival outcomes within defined selection windows.

## Methods

### Yeast strains and plasmid construction

The yeast strains, plasmids, and oligonucleotides used for plasmid construction and gene modification are listed in Supplementary File 1. All yeast strains used in this study are derivatives of *Saccharomyces cerevisiae* W303-1A (MATa ade2-1 can1-100 his3-11,15 leu2-3,112 trp1-1 ura3-1). We amplified the coding sequences of *URA3* and *TRP1* and placed them under the control of the native *TIM50* promoter (a moderate constitutive promoter) and the *TIM50* 3′ UTR. To facilitate MS2-mediated localization, a sequence encoding 12 copies of the MS2 coat protein binding hairpin (MS2 tag) was fused to the 3′ UTR of each gene (before the terminator). A short flexible linker separated the gene’s stop codon and the MS2 tag array. The *URA3* and *TRP1* constructs (with promoter, ORF, MS2 tag, terminator) were cloned into integrative plasmid backbones derived from pRS403 (*HIS3* integrating vectors). These plasmids, named PYL001 (*URA3*-MS2) and PYL002 (*TRP1*-MS2), were linearized at the *HIS3* locus and transformed into yeast to target insertion into the his3-11,15 locus (the W303-1A his3 allele). Standard lithium acetate transformation was used. Our chemical transformation integration at the location of the mutated *HIS3* gene can randomly generate strains with different copy numbers (Figure 1C, right). Correct integration and copy number were verified by PCR MS2 sequence from the yeast genome using MS2-F and MS2-R primers, followed by electrophoresis^11^. The TOM70–MCP fusion gene was cloned into an integrative plasmid pRG205, which integrates 1 copy of the plasmid into the cells, from the cells, integrated MCP sequence (in lab stocks). The construct (TTP235 or TTP242, Table 1) was integrated into the *LEU2* locus of strains already carrying the *URA3*-MS2 or *TRP1*-MS2 constructs.

### Media and growth conditions

For selection of *URA3* marker, we used SC medium lacking uracil (SC–Ura) to select for *URA3*^+^ cells, and supplemented with 5-fluoroorotic acid (US Biological) for counter-selection. 5-FOA plates were made with yeast nitrogen base (YNB) complete minus uracil, 2% glucose, and 5-FOA at concentrations ranging from 0.3 to 1.0 g/l, plus 50 mg/l uracil (a small amount of uracil is commonly added to 5-FOA plates to reduce background and allow slight growth of Ura3^+^ cells before they convert 5-FOA to toxin)^6^. For *TRP1* selection, SC medium lacking tryptophan (SC– Trp) was used to select *TRP1*^+^ transformants. 5-FAA plates were prepared as described by Toyn *et al*. (2000)^3^ with some modifications: briefly, YNB complete minus tryptophan with 5% glucose and agar was supplemented with 5-fluoroanthranilic acid (Toronto Research Chemicals) at 100 mg/L for initial tests. We found that including a minute amount of tryptophan (5–10 µg/mL) in 5-FAA plates improved the differentiation of phenotypes, likely by allowing basal growth of cells that completely lack Trp1p, so that partial Trp1p activity can be toxic in comparison. For spot assays, saturated overnight cultures were adjusted to OD_600_ ∼1.0, then ten-fold serially diluted in sterile water and spotted (5 µL) onto plates. Plates were photographed after 2–4 days of incubation at 30°C. Each experiment was performed at least in triplicate to ensure reproducibility.

## Data and materials availability

All data supporting the findings of this study are available within the paper and its supplementary information. Yeast strains, plasmids, and oligonucleotides used in the study are provided in Supplementary File 1. Further information and requests for raw data and reagents should be directed to and will be fulfilled by the lead contact, T.T.

## Supplemental information

Supplementary File 1 | List of yeast strains, plasmids, and oligonucleotides.

## Acknowledgments

We thank members of the Tsuboi laboratory for helpful discussions and feedback on the manuscript. This work was supported in part by the Key Research and Development Program of the Ministry of Science and Technology (2024YFE0102700, 2023YFA0914303), the Shenzhen Science and Technology Innovation Commission (WDZC20220811144737001), startup fund OD2021031C, Interdisciplinary Research and Innovation Fund JC2022008, and Overseas Research Cooperation Fund HW2024009 from Tsinghua SIGS (to T.T).

## Author contributions

T.T. conceived and designed the project. Y.L. performed the experiments and analyzed image data. Y.L. and T.T. wrote the manuscript.

## Competing interests

The authors declare no competing interests.

## Notes

### Competing Interest Statement

The authors have declared no competing interest.

## References

1. Galletta, B.J., and Rusan, N.M. (2015). A yeast two-hybrid approach for probing protein-protein interactions at the centrosome. Methods Cell Biol 129, 251–277. 10.1016/bs.mcb.2015.03.012.

2. Boeke, J.D., Trueheart, J., Natsoulis, G., and Fink, G.R. (1987). 5-Fluoroorotic acid as a selective agent in yeast molecular genetics. Methods Enzymol 154, 164–175. 10.1016/0076-6879(87)54076-9.

3. Toyn, J.H., Gunyuzlu, P.L., White, W.H., Thompson, L.A., and Hollis, G.F. (2000). A counterselection for the tryptophan pathway in yeast: 5-fluoroanthranilic acid resistance. Yeast 16, 553–560. 10.1002/(sici)1097-0061(200004)16:6<553::Aid-yea554>3.0.Co;2-7.

4. Tsuboi, T., Viana, M.P., Xu, F., Yu, J., Chanchani, R., Arceo, X.G., Tutucci, E., Choi, J., Chen, Y.S., Singer, R.H., et al. (2020). Mitochondrial volume fraction and translation speed impact mRNA localization and production of nuclear-encoded mitochondrial proteins. Elife 9, 1–24.

5. Miura, F., Kawaguchi, N., Yoshida, M., Uematsu, C., Kito, K., Sakaki, Y., and Ito, T. (2008). Absolute quantification of the budding yeast transcriptome by means of competitive PCR between genomic and complementary DNAs. BMC Genomics 9, 1–14. 10.1186/1471-2164-9-574.

6. Ko, N., Nishihama, R., and Pringle, J. (2008). Control of 5-FOA and 5-FU resistance bySaccharomyces cerevisiae YJL055W. Yeast 25, 155–160. 10.1002/yea.1554.

7. Lacroute, F. (1968). Regulation of pyrimidine biosynthesis in Saccharomyces cerevisiae. J Bacteriol 95. 10.1128/jb.95.3.824-832.1968.

8. Miozzari, G., Niederberger, P., and Huetter, R. (1978). Tryptophan biosynthesis in Saccharomyces cerevisiae: control of the flux through the pathway. J Bacteriol 134. 10.1128/jb.134.1.48-59.1978.

9. García-Rodríguez, L.J., Gay, A.C., and Pon, L.A. (2007). Puf3p, a Pumilio family RNA binding protein, localizes to mitochondria and regulates mitochondrial biogenesis and motility in budding yeast. J Cell Biol 176, 197–207. 10.1083/jcb.200606054.

10. Eliyahu, E., Pnueli, L., Melamed, D., Scherrer, T., Gerber, A.P., Pines, O., Rapaport, D., and Arava, Y. (2010). Tom20 Mediates Localization of mRNAs to Mitochondria in a Translation-Dependent Manner. Mol Cell Biol 30, 284–294. 10.1128/MCB.00651-09.

11. Lõoke, M., Kristjuhan, K., and Kristjuhan, A. (2011). Extraction of genomic DNA from yeasts for PCR-based applications. Biotechniques 50, 325–328. 10.2144/000113672.

