## Supplemental File 1 for "Expanding tunable selection in yeast using auxotrophic markers *URA3* and *TRP1*"

### Supplementary File 1. Yeast strains, plasmids, and primers used in this study

| Name | Description | Source |
|---|---|---|
| **Strains:** | | |
| W303-1A (parent) | MATa ade2-1 can1-100 his3-11 leu2-3 trp1-1 ura3-1 | Lab stock |
| YYL001 | W303-1A *TIM50p-Flag-URA3::12×MS2 tag::TIM50 3'UTR* (1 copy) | This study |
| YYL002 | Same as YYL001, with 2 copies of *URA3* construct | This study |
| YYL003 | Same as YYL001, with 4 copies of *URA3* construct | This study |
| YYL004 | W303-1A *TIM50p-Flag-TRP1::12×MS2 tag::TIM50 3'UTR* (1 copy) | This study |
| YYL005 | Same as YYL004, with 2 copies of *TRP1* construct | This study |
| YYL006 | Same as YYL004, with 4 copies of *TRP1* construct | This study |
| YYL007 | YYL001 TOM70-MCP | This study |
| YYL008 | YYL002 TOM70-MCP | This study |
| YYL009 | YYL003 TOM70-MCP | This study |
| YYL010 | W303-1A TOM70-MCP | This study |
| YYL011 | YYL004 TOM70-MCP | This study |
| **Plasmids:** | | |
| pRS406 | Integrative *URA3*-marked empty vector | Lab stock |
| pRS404 | Integrative *TRP1*-marked empty vector | Lab stock |
| TTP158 | pRS403*TIM50p-flagyoGFP-TIM50ter-MS2tag* | This study |
| TTP235 | pRG205*TOM70p-TOM70-MCP-TOM70ter* | This study |
| PYL001 | pRS403*TIM50p-flagURA3-TIM50ter-MS2tag* | This study |
| PYL002 | pRS403*TIM50p-flagTRP1-TIM50ter-MS2tag* | This study |
| **Primers:** | | |
| TIM50-URA3-F | GGACTACAAGGACGACGATGACAAGATGTCGAAAGCTACATATAAGGAACGTGC | |
| TIM50-URA3-R | GGTTGTCGACCTGCAGCGTTTTAGTTTTGCTGGCCGCATCTTCTCAAATATG | |
| TIM50-TRP1-F | GGACTACAAGGACGACGATGACAAGATGTCTGTTATTAATTTCACAGG | |
| TIM50-TRP1-R | TATTAAGGGTTGTCGACCTGCAGCGTTCTATTTCTTAGCATTTTTGACGAAATTTGC | |
| PYL1-Vector-F1 | TAAAACGCTGCAGGTCGACAACC | |
| PYL2-Vector-F2 | TAAAACGCTGCAGGTCGACAACCCTTAATATAACTTCG | |
| PYL2-Vector-R2 | CTTGTCATCGTCGTCCTTGTAGTCCATTGCAAG | |
| MS2-F | GCTATACGAAGTTATTAGGTGATATCA | |
| MS2-R | GGGTTTACATGAAAAGCATAGGC | |
